# Evaluating discrepancies in dimensionality reduction for time-series single-cell RNA-sequencing data

**DOI:** 10.1101/2025.02.06.636822

**Authors:** Maren Hackenberg, Laia Canal Guitart, Rolf Backofen, Harald Binder

**Affiliations:** Institute of Medical Biometry and Statistics, Faculty of Medicine and Medical Center, University of Freiburg, Germany; Freiburg Center for Data Analysis and Modeling, University of Freiburg, Germany; Bioinformatics Group, Department of Computer Science, University of Freiburg, Germany; Centre for Integrative Biological Signaling Studies (CIBSS), University of Freiburg, Germany

**Keywords:** single-cell RNA-sequencing, time-series data, dimensionality reduction, evaluation, synthetic data, deep learning

## Abstract

There are various dimensionality reduction techniques for visually inspecting dynamical patterns in time-series single-cell RNA-sequencing (scRNA-seq) data. However, the lack of one-to-one correspondence between cells across time points makes it difficult to uniquely uncover temporal structure in a low-dimensional manifold. The use of different techniques may thus lead to discrepancies in the representation of dynamical patterns. However, The extent of these discrepancies remains unclear. To investigate this, we propose an approach for reasoning about such discrepancies based on synthetic time-series scRNA-seq data generated by variational autoencoders. The synthetic dynamical patterns induced in a low-dimensional manifold reflect biologically plausible temporal patterns, such as dividing cell clusters during a differentiation process. We consider manifolds from different dimensionality reduction techniques, such as principal component analysis (PCA), t-distributed stochastic neighbor embedding (t-SNE), uniform manifold approximation and projection (UMAP) and single-cell variational inference (scVI). We illustrate how the proposed approach allows for reasoning about to what extent low-dimensional manifolds, obtained from different techniques, can capture different dynamical patterns. The results indicate that these techniques may not reliably represent dynamics when used in isolation. Thus, the proposed synthetic dynamical pattern approach provides a foundation for guiding future methods development to detect complex patterns in time-series scRNA-seq data.

## 1 Introduction

Dimensionality reduction is an essential component of any exploratory single-cell RNA sequencing (scRNA-seq) analysis workflow to reveal the underlying patterns of cell heterogeneity, and many computational approaches have been proposed [1, 2]. Among the most popular techniques are t-stochastic neighborhood embedding (t-SNE) [3] and uniform manifold approximation and projection (UMAP) [4, 5]. They come in many recently developed variants, such as parametric versions [6, 7, 8], versions with explainability components [9] or versions for longitudinal structure [10]. Other commonly used approaches include linear techniques based on principal component analysis (PCA) [11]. While PCA, t-SNE, and UMAP are often used for visualization in two or three dimensions, there are also deep learning–based methods that support further downstream tasks beyond visualization, such as batch correction, noise reduction, imputation or synthetic data generation [12, 13, 14]. The latter often use an autoencoder architecture, where a low-dimensional representation is learned in an unsupervised way via artificial neural networks, e.g., in the single-cell variational inference (scVI) framework [12, 15], or [16, 17].

These dimensionality reduction techniques are also used for more complex settings, in particular with experimental designs that comprise multiple time points to provide insights into dynamic biological processes such as differentiation, proliferation, or response to stimuli [e.g., 18, 19, 20]. Since the sequencing protocol is usually destructive, there is no one-to-one correspondence between cells at different time points, making it difficult to identify cellular trajectories [21]. Although there are approaches to computationally trace cell populations over time [e.g. 22, 23, 24, 25], these often tend to be rather complex and specific to the scenario for which they have been developed. In practice, researchers often resort to simultaneous dimensionality reduction of the data from all time points using PCA, t-SNE or UMAP, with subsequent visualization and comparison between time points [26, 27, 28, 29].

While these methods are known to effectively cluster cells based on similarities in their gene expression profiles [30], they are not explicitly designed to capture biological dynamics [31], i.e., they are used under the implicit assumption that they capture the underlying lower-dimensional manifold in which the dynamics occur [32]. As this assumption is not explicitly enforced, the obtained representation does not necessarily have to coincide with the one that best reflects the underlying structure. This means that researchers cannot reliably use the spatial arrangement of clusters seen in visualizations across time points to infer how cells transition between states because continuous relationships between cells in high-dimensional gene expression space might be distorted or lost [33, 34]. Indeed, an increasing number of results suggest that such representations might be misleading [35, 36, 34, 37]. However, the extent of such potential problems has not been investigated systematically so far. We therefore propose an approach for reasoning about discrepancies in dimensionality reduction representations of temporal patterns.

Two main strategies are used for evaluating dimensionality reduction techniques for scRNA-seq data, namely, using real data or simulated data. For settings without temporal structure, current benchmarks almost exclusively consider clustering performance with real data, such as for cell type discovery, i.e., structures where the ground truth is known (e.g., [38, 39, 40]). Unfortunately, such ground-truth information is typically not available in the analysis of time-series data. While there are some approaches for simulating scRNA-seq data with temporal structure, such as the splatter R package [41], these typically do not allow for introducing more complex developmental patterns. In particular, there is no consideration of the original manifold in which the patterns occur, making it difficult to assess whether a dimensionality reduction approach has identified an appropriate representation. Therefore, we propose a novel approach in which we extract manifolds from real data and directly induce temporal patterns in these.

We consider deep generative approaches for obtaining manifolds and generating synthetic data, which are a promising tool for obtaining realistic synthetic single-cell RNA-seq data [14, 42]. Specifically, we suggest using a variational autoencoder (VAE) [43] for generating synthetic time-series data based on a snapshot scRNA-seq dataset. We propose an approach for representing low-dimensional manifolds from different techniques via VAEs, specifically from PCA, t-SNE, UMAP, and scVI. In these manifolds, we use vector fields that describe different plausible temporal patterns, such as dividing cell clusters during a differentiation process. Applying these vector fields to each manifold, we can simulate a dynamic process directly within the low-dimensional space, reflecting biologically meaningful cellular transitions. The VAE then can map back from the transformed manifold to gene expression space to create benchmark datasets with different known underlying structures (i.e., different temporal patterns introduced in different underlying manifolds).

We illustrate the capabilities of the proposed approach to investigate to what extent low-dimensional manifolds, obtained from different techniques, can capture different dynamical patterns in an evaluation of various dimensionality reduction techniques. As a starting point, we describe the typical strategy currently used for dimensionality reduction and visualization of temporal scRNA-seq data before giving a brief overview of specific dimensionality reduction techniques, namely PCA, t-SNE, and UMAP. These are subsequently investigated and compared with several dynamic patterns induced by the proposed synthetic data approach, followed by a discussion of the discrepancies and consequences for data analysis strategy and future methods development.

We provide an implementation of our approach, including tutorial notebooks on Github at https://github.com/laia-cg/scManifoldDynamics.

## 2 Materials and methods

### 2.1 Illustration of a typical strategy for dimensionality reduction on a timeseries scRNA-seq dataset

When visually analyzing temporal patterns in time-series scRNA-seq data using dimensionality reduction, thedrop techniques are typically applied to the joint dataset of all cells from all time points. This embedding projects the entire dataset into a common low-dimensional space. For visualization, the embedded representation is then split by time point, i.e., for each time point, a separate plot is created, showing only the cells captured at that time point in the joint representation. This allows for observing the distribution of cells at each time point separately and for comparing the locations of cell type clusters across time points to trace developmental patterns visually.

Figure 1 illustrates this procedure, applying PCA (Panel A), t-SNE (Panel B), UMAP (Panel C), and scVI (Panel D) in turn on a time-series dataset of human embryonic stem cells grown as embryoid bodies from four time points spanning 27 days during differentiation [32]. In this example, while PCA does not resolve most of the local cluster structure in the data, some structures are captured to some extent by all of the other three methods, e.g., a transition from the cells indicated in yellow via the subgroup indicated in brown, and a subgroup indicated in grey to the cells shown in blue. However, the general visual impression of the emerging temporal pattern is rather distinct, indicating that different methods come to a relatively different representation of the underlying biological development process. From the plot alone and in the absence of ground-truth information, it is not clear, however, whether these discrepancies are merely due to different properties of each algorithm and optimization procedure, e.g., UMAP generally produces sharper clusters than scVI, where the latent representation is modeled as a Gaussian distribution, or whether there are qualitative differences in the perspective of different approaches on the underlying dynamics.

**Figure 1:**
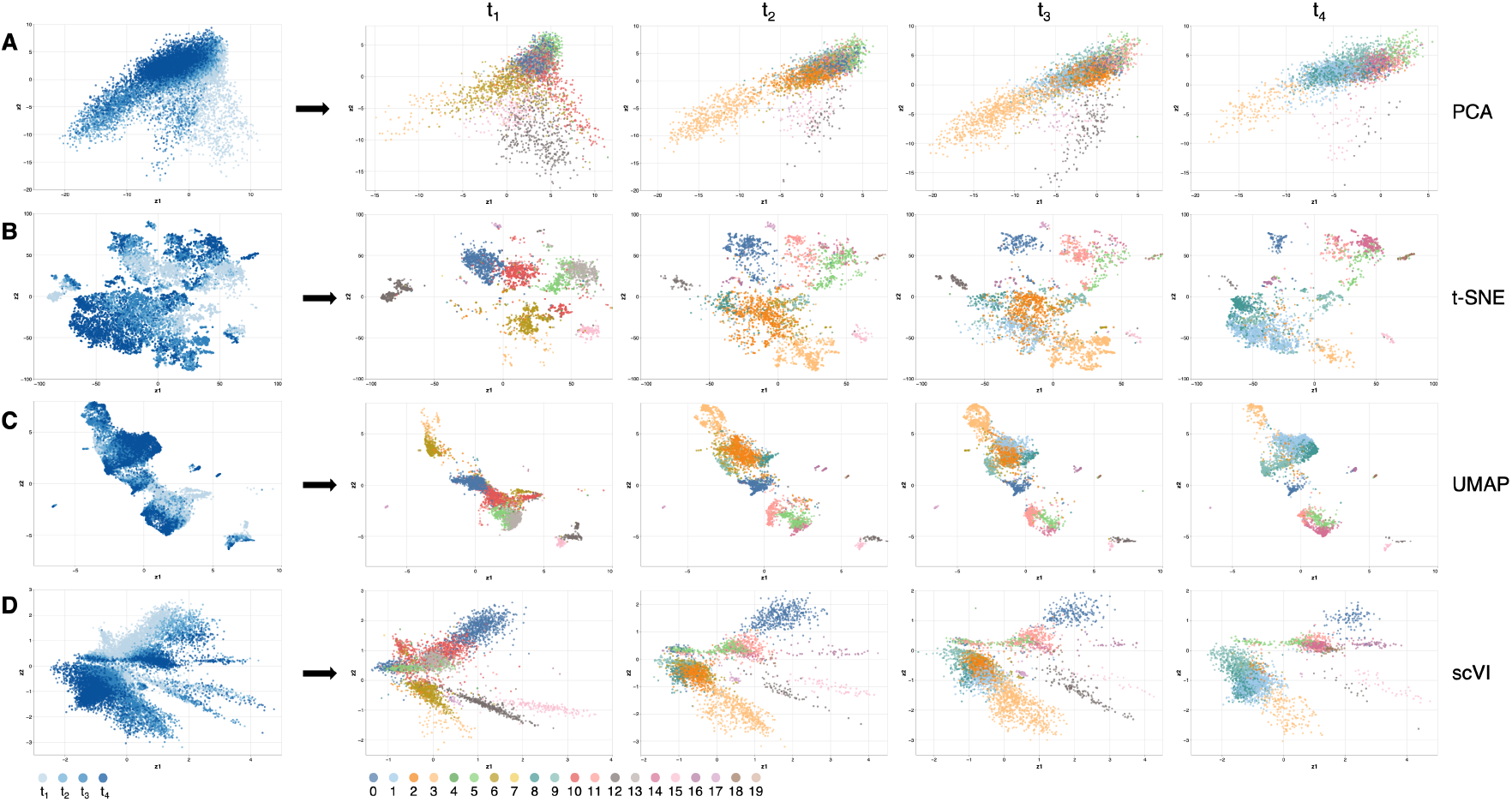
Different dimensionality reduction methods applied to the Embryoid body dataset [32]. The different rows correspond to different dimensionality reduction techniques applied to the dataset: The first row shows dimensionality reduction using PCA, the second row shows representations obtained using t-SNE, the third row corresponds to UMAP, and the last row to the scVI representation. The first column displays the entire data set, color-coded by time point, with the progression of time represented by a range of shades from light blue to dark blue. The subsequent columns represent only the cells from the four selected time points, color-coded by Louvain clusters. Clustering follows the original authors’ specifications. For each method, we calculated a joint dimensionality reduction, using the cells from all time points and split it by time point for visualization.

### 2.2 Dimensionality reduction techniques as manifold learning tools to reveal cellular dynamics

Using dimensionality reduction to visualize temporal structure in scRNA-seq data assumes that the underlying cellular state space is a lower-dimensional manifold, where cells transition smoothly between states. Waddington’s landscape metaphor of cellular development [44], though limited given today’s knowledge on cellular differentiation [45, 46], illustrates this idea. Differentiation is conceptualized as the process of a marble (corresponding to a cell) rolling down a landscape of hills and valleys (the cellular state space manifold), determining the cell’s fate. Time-series scRNA-seq data corresponds to snapshots of this process at different stages.

Any dimensionality reduction approach generates a possible version of this underlying, intrinsically unobserved, cellular state space. When visualizing this manifold, similar cells typically cluster together [31]. In stable, fully differentiated cell populations, using dimensionality reduction for visualization may be sufficient to understand their organization. However, for dynamic processes, dimensionality reduction techniques should not only place similar cells close to each other but also reflect developmental trajectories.

We focus on four commonly used dimensionality reduction techniques, covering a range of complexity and different prototypical mechanisms [47]. Specifically, we consider PCA as a deterministic matrix factorization-based linear method, nonlinear and probabilistic neighbor-based algorithmic approaches like t-SNE and UMAP, and scVI as a nonlinear, probabilistic method based on neural networks. These different characteristics of the approaches are reflected in their underlying objective when optimizing an embedding – finding the rotation and projection that capture maximum variance, embedding a graph or a distribution over nearest neighbors, or minimizing a reconstruction loss (Figure 2 **A**). All of them work unsupervised and optimize data-intrinsic properties without allowing to incorporate structural information, e.g., on temporal dynamics, in their original (and widely used) version.

**Figure 2:**
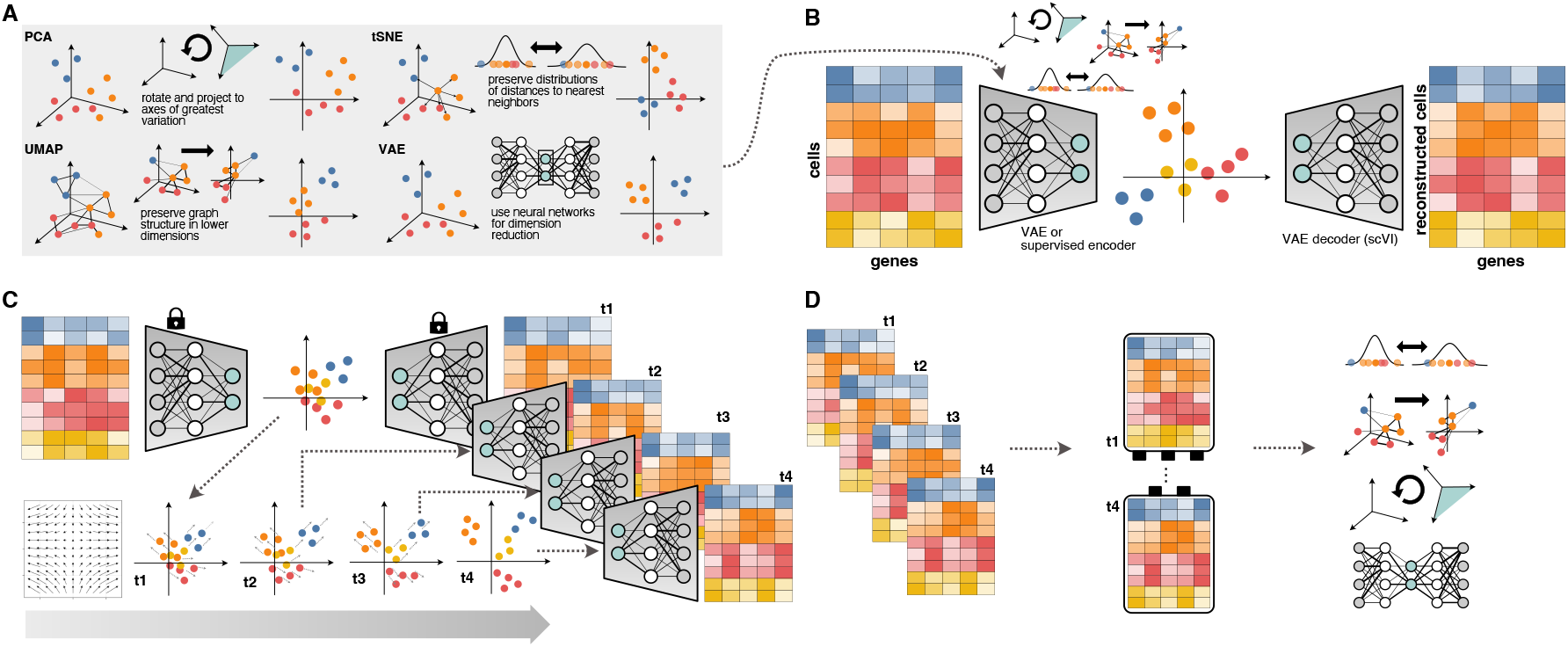
Schematic illustration of the proposed approach to generate synthetic data with dynamic temporal structure for comparing dimensionality reduction approaches. **A:** Different popular dimensionality reduction approaches for scRNA-seq data. **B:** We train a (modified) VAE on a snapshot scRNA-seq dataset to obtain (1) a two-dimensional representation, which can optionally mimic a t-SNE, UMAP, or PCA manifold via supervised training, and (2) a matching scVI decoder, which can generate single-cell data based on the respective manifold. **C:** Next, we introduce artificial dynamic patterns in the obtained manifold by applying vector fields and use the trained VAE to generate corresponding high-dimensional data for each synthetic time point. **D:** We concatenate the datasets to create a synthetic time-series dataset, apply different dimensionality reduction techniques and compare.

#### 2.2.1 Principal component analysis

Principal component analysis (PCA) is a linear technique for dimensionality reduction based on matrix factorization by singular value decomposition (SVD) [48]. Specifically, SVD provides a rotation of the original data matrix such that the variation within the data happens along the new coordinate axes, called the principal components (PCs). By projecting onto the first two PCs, a dimensionality reduction in 2D is obtained, which retains as much of the original variation in the data as possible.

PCA is a linear, global approach which does not capture fine-grained local structures or nonlinearities [49]. Further, it may be sensitive to outliers and the scaling of the data [50, 51].

#### 2.2.2 t-distributed stochastic neighbor embedding

To capture more complex, nonlinear patterns in the data, t-distributed stochastic neighbor embedding (t-SNE) [3] has emerged as a popular tool for single-cell transcriptomics [1, 9]. The approach builds a probability distribution over pairwise distances of cells in the high-dimensional gene expression space, such that for each cell, other cells with similar transcriptomic profiles are assigned a higher probability. t-SNE then embeds points into a two-dimensional space by defining a corresponding distribution over the low-dimensional distances and minimizing the Kullback-Leibler divergence between the two distributions via gradient descent. The resulting embedding preserves local neighbor relations between similar cells, i.e., relative distances. t-SNE is thus designed to focus on local rather than global structure of the data. While it provides finely resolved cell clusters, the global distances between clusters cannot be meaningfully interpreted [52]. The approach can be sensitive to the choice of hyperparameters and the initialization of the algorithm [1]. It is computationally rather intensive and thus does not scale well to large datasets. t-SNE is an inherently nonparametric technique. As the embedding is optimized for each cell in relation to its neighbors, for projecting new cells onto a given embedding, the embedding has to be re-computed based on the new neighbor relations. To circumvent this, parametric alternatives employ neural networks to learn a functional mapping that approximates the t-SNE embedding [53, 7, 54].

#### 2.2.3 Uniform manifold approximation and projection

Similar to t-SNE, the uniform manifold approximation and projection (UMAP) algorithm is a nonlinear technique for dimensionality reduction that embeds points in a low-dimensional space, preserving local neighborhood information. Specifically, it constructs a kNN-based neighbor graph that is subsequently embedded into a lower-dimensional space. UMAP is theoretically grounded in Riemannian geometry as it is based on the assumption that the data is uniformly distributed on a locally connected Riemannian manifold and that the Riemannian metric is (approximately) locally constant [4]. While producing visually similar representations to t-SNE, its main advantages over t-SNE are that it can better capture global relations to some extent [4, 55]. This is achieved by the constructing fuzzy graphs that can also capture long-distance relationships beyond immediate local neighbor information and thus better approximate the overall topology. Further, UMAP is more scalable due to a more efficient optimization algorithm. However, similar to t-SNE, it is sensitive to hyperparameters and evidence suggests that it effectively binarizes similarity information [52, 36]. Like t-SNE, it is nonparametric and requires re-optimization to embed new data. A parametric version using neural networks has been proposed [8].

#### 2.2.4 Single-cell variational inference

An autoencoder consists of an encoder and decoder neural network that map data to a lower-dimensional latent space and back (Figure 2, A). The neural network parameters are optimized by minimizing the reconstruction error between the original input and the reconstructed output, such that the model learns to compress the main characteristics of the data in the lower-dimensional space. This approach has been successfully employed for dimensionality reduction of scRNA-seq data [17]. The variational autoencoder (VAE) extends the autoencoder to a probabilistic version [43, 56] by modeling the latent space as a random variable, using variational inference [57] to approximate the conditional probability distributions of the latent variable given the data and vice versa in the encoder and decoder. Training maximizes a lower bound on the true data likelihood called the evidence lower bound (ELBO), such that the model learns to approximate the underlying high-dimensional data distributions, enabling subsequent synthetic data generation. For scRNA-seq, the scVI model adapts VAEs to negative binomial distributions while accounting for library size and batch effects [12, 58, 59, 60]. Synthetic scRNA-seq data from such models can be leveraged, e.g., for investigating dominant patterns in scRNA-seq data [**61**] or experimental planning [14, 42]. The low-dimensional representation can be used for downstream tasks such as classification or clustering [12, 15]. While such analyses are typically based on a richer representation with more latent dimensions, here we focus on the ability of scVI to visually capture temporal patterns in comparison to PCA, t-SNE, and UMAP and therefore use a two-dimensional latent space.

#### 2.3 Generating synthetic time-series scRNA-seq data, using different manifolds for introducing temporal structure

Our approach for reasoning about dynamical patterns in different manifolds is based on synthetic timeseries scRNA-seq data, to enable access to ground-truth information. For synthetic data generation, we use a VAE approach based on the scVI framework [12, 15]. We train a VAE on a real snapshot scRNA-seq dataset, i.e., without temporal structure, and subsequently introduce a synthetic temporal pattern via vector field dynamics, as explained below. To explore dynamics in different manifolds, we adapt scVI by introducing a supervised encoder, such that the latent representation is trained to align with a given t-SNE, UMAP or PCA manifold (Figure 2 B). We chose this VAE-based approach, because t-SNE and UMAP are nonparametric, i.e., do not represent the embedding into low-dimensional space as an explicit function and cannot generate new data points, whereas the VAE is a parametric approach that is inherently generative, i.e., allows for creating synthetic data.

To train the adapted scVI model with a supervised encoder, we add the mean squared error between the VAE latent representation and the coordinates of a precomputed t-SNE, UMAP, or PCA embedding to the loss function. The trained encoder then acts as a surrogate to parameterize the embedding function of t-SNE, PCA, or UMAP. Given a snapshot dataset, we thus compute t-SNE, PCA, and UMAP manifolds, and train VAEs with supervised encoders for each, alongside a standard scVI model for a pure VAE manifold. For each approach, we use two-dimensional manifolds for visualization.

We select one dimensionality reduction approach as the “ground-truth” manifold for simulating temporal structure, and use the corresponding supervised encoder VAE. Then, we introduce temporal dynamics by transforming the two-dimensional representation using a vector field (Figure 2 C). This geometric perspective allows for applying biologically meaningful transformations to the geometry of the cell manifold as a whole rather than specifying individual cellular trajectories, aligning with the landscape metaphor [62, 63].

Formally, a vector field **F** : ℝ^*d*^ → ℝ^*d*^ on a set 𝒟 ⊂ ℝ^*d*^ assigns to each point **x** ∈ 𝒟 a vector **F**(**x**) = (*F*_1_(**x**), …, *F*_*d*_(**x**)). For our applications in ℝ^2^, the vector field is associated with a dynamical system governed by the differential equations:

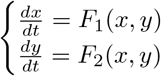

This formulation helps in defining transformations *P* and *Q* which can mimic the behavior of the dynamical system without needing explicit solutions to the differential equations.

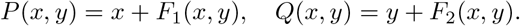

In practice, *P* and *Q* are each defined as a series of transformations, each dependent on a set of parameters *θ* that tailor each transformation to the desired behavior. As a result, the components *P* and *Q* transform any point **p** = (*x, y*) in the domain 𝒟 as

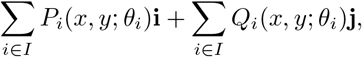

where **i** and **j** are the unit vectors in the *x* and *y* directions, respectively and *I* is the set of indices for the transformations.

The additive structure allows for stacking various transformations to manipulate the vector field **F**, making the design visually intuitive and accessible for users. We provide a detailed guide on how to design customized transformations based on this approach, available as a tutorial notebook in our Github repository. While our framework allows for freely customizable transformations, we also provide predefined vector fields for common use cases as ready-to-use solutions in our implementation. The vector fields are designed to correspond to meaningful and biologically plausible developmental patterns, e.g., dividing and spreading cell groups, as illustrated in Figure 2 **C**.

We apply the vector field dynamics to the two-dimensional representation of the snapshot dataset and iterate this process over a fixed number of time steps. At each step we also add a small random offset to each cell’s representation, to account for the stochasticity inherent in cellular evolution and potential measurement noise. The equations describing the dynamics are tailored specifically for each dataset and incorporate available cell type annotations.

To generate corresponding high-dimensional data, we pass the transformed latent space from each artificial time point to the VAE decoder, obtaining parameters of a high-dimensional negative binomial distribution for sampling. This generates synthetic high-dimensional time-series data from a real snapshot dataset with artificially introduced temporal structure, specified as a used-defined vector field, in a chosen low-dimensional manifold. Since both the manifold in which the dynamics happen and the dynamical process itself are known, this approach provides a controlled benchmark with a known ground truth.

Finally, we concatenate the generated data across time points to obtain a complete time-series dataset, on which we apply all dimensionality reduction approaches (Figure 2 D). Knowing the original manifold and the true underlying pattern then enables us to reason about discrepancies in their representations, and investigate the sensitivity of the different approaches with respect to the manifold in which the dynamics happen, and to the dynamical pattern itself.

#### 3 Results

To illustrate our approach across different contexts, we have applied it on two snapshot scRNA-seq datasets, a dataset of 8000 peripheral blood mononuclear cells (PBMCs) [64], referred to as the *PBMC data*, and a dataset of 3000 mouse brain cells [65], which we refer to as the *Zeisel data*. These two datasets have been used as the initial basis at the first time point for applying our method to generate the high-dimensional synthetic time-series datasets.

For each dataset, we have compared different hyperparameter configurations of t-SNE, UMAP and scVI, and selected the parameters that provided the visually most informative representation, reflecting the experience of a domain expert user without in-depth programming expertise. Even if there is some combination of pre-processing and hyperparameters that might lead to “better” capturing a specific pattern, it would be impossible to identify this without access to the ground truth. We have not found any systematic structure in hyperparameter configurations that would in general allow for better seeing temporal patterns. All plots are colored according to original cell type annotation from the first timepoint. As this annotation is no longer consistent when clusters experience differentiating processes, we refer to the clusters by their color annotation and not cell type label.

Figure 3 shows an exemplary result on the PBMC data. In this case, the blue cluster differentiates, creating two new separate clusters: one exhibits increasingly similar gene expression profiles to those of the yellow cluster, whereas the other one shows a growing resemblance to the profiles of the pink and red cell types. We have taken the t-SNE and UMAP representations based on the original data as first time points, and transformed the cell clusters according to the described pattern in both spaces (Figure 3 **A** and Figure 3 **C** respectively). From each of these artificial time points, we have then generated corresponding high-dimensional gene expression datasets, and applied t-SNE, UMAP, PCA and scVI on the resulting dataset. Our design induces temporal patterns in different manifolds, allowing us to evaluate how the same transformation is captured by different methods across these manifolds. As expected, when the differentiation process occurs in the t-SNE manifold, t-SNE effectively depicts the transformation (Figure 3 **B**), likely because the method is operating in its own representation space. However, when the same transformation takes place in the UMAP manifold, t-SNE can still capture the differentiation, but it is visually less evident (Figure 3 **D**).

**Figure 3:**
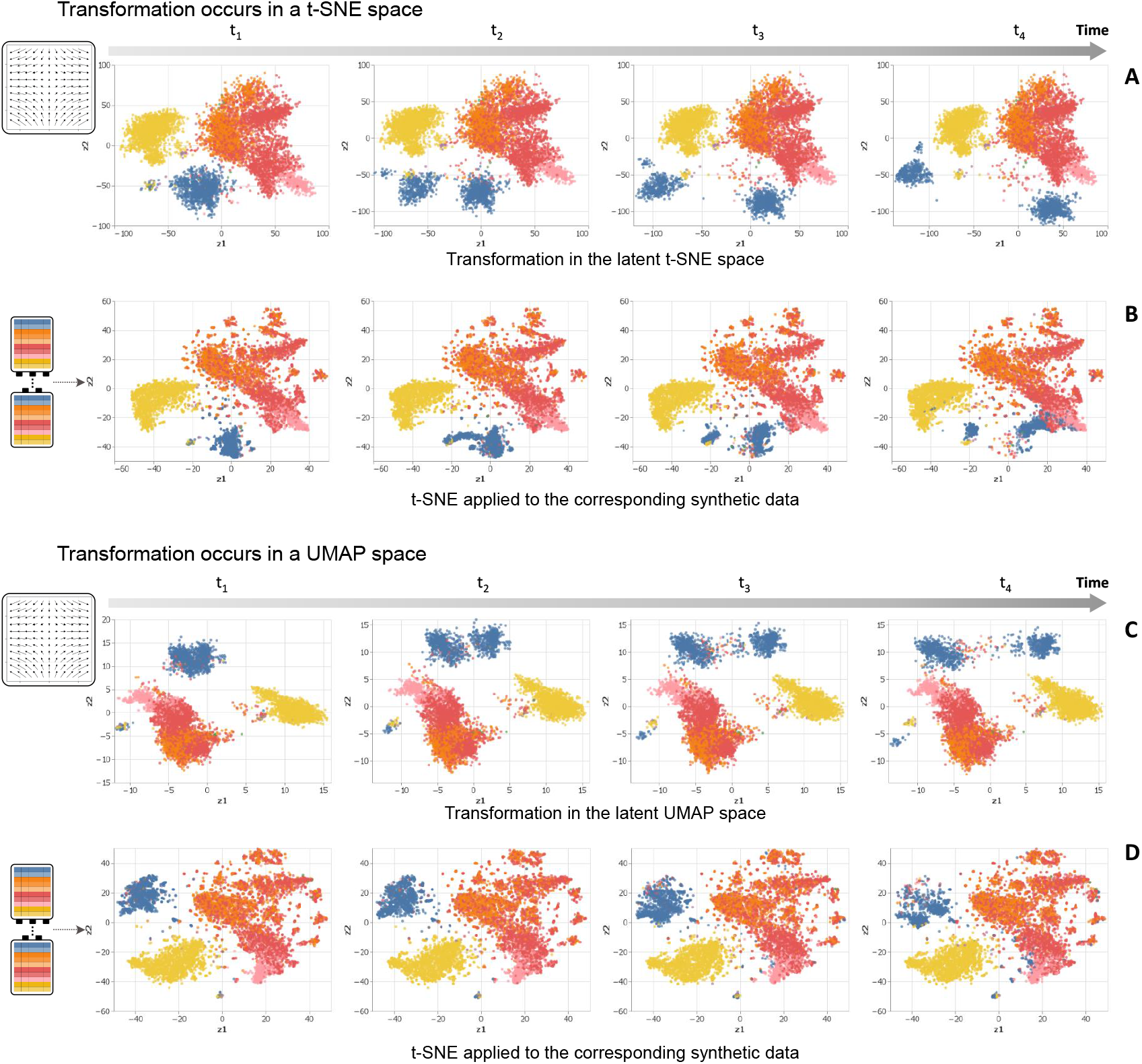
Illustrative example of a comparative analysis of how dimensionality reduction methods capture similar artificially induced temporal structures across different manifolds in scRNA-seq data. **(A**,**C)**: Latent representations from the supervised VAE, trained to match the t-SNE (A) and UMAP (C) embeddings of the PBMC dataset (leftmost panel), used as initial time point (t1), and transformed to induce a visual temporal pattern corresponding to the differentiation of the blue cell type cluster (panels t2-t4). **(B**,**D)**: t-SNE applied to the high-dimensional datasets generated by decoding the transformed latent representations in the t-SNE (A) and UMAP (C) manifolds using the supervised VAE. This comparison illustrates how the same transformation is captured differently by t-SNE across these manifolds. Colors correspond to manually annotated cell types based on the original time point (t1).

This inconsistency is observed across various combinations of methods and manifolds, where certain methods perform better in specific manifolds and for particular transformations, but no single method dominates across all scenarios. For example we can observe in Figure 4 how PCA enables us to see two cell clusters evolving in different directions in terms of their gene expression profiles within a UMAP manifold (Figure 4 **B**), but the same type of transformations is not perceptible anymore when it involves different clusters and directions (Figure 4 **D**).

**Figure 4:**
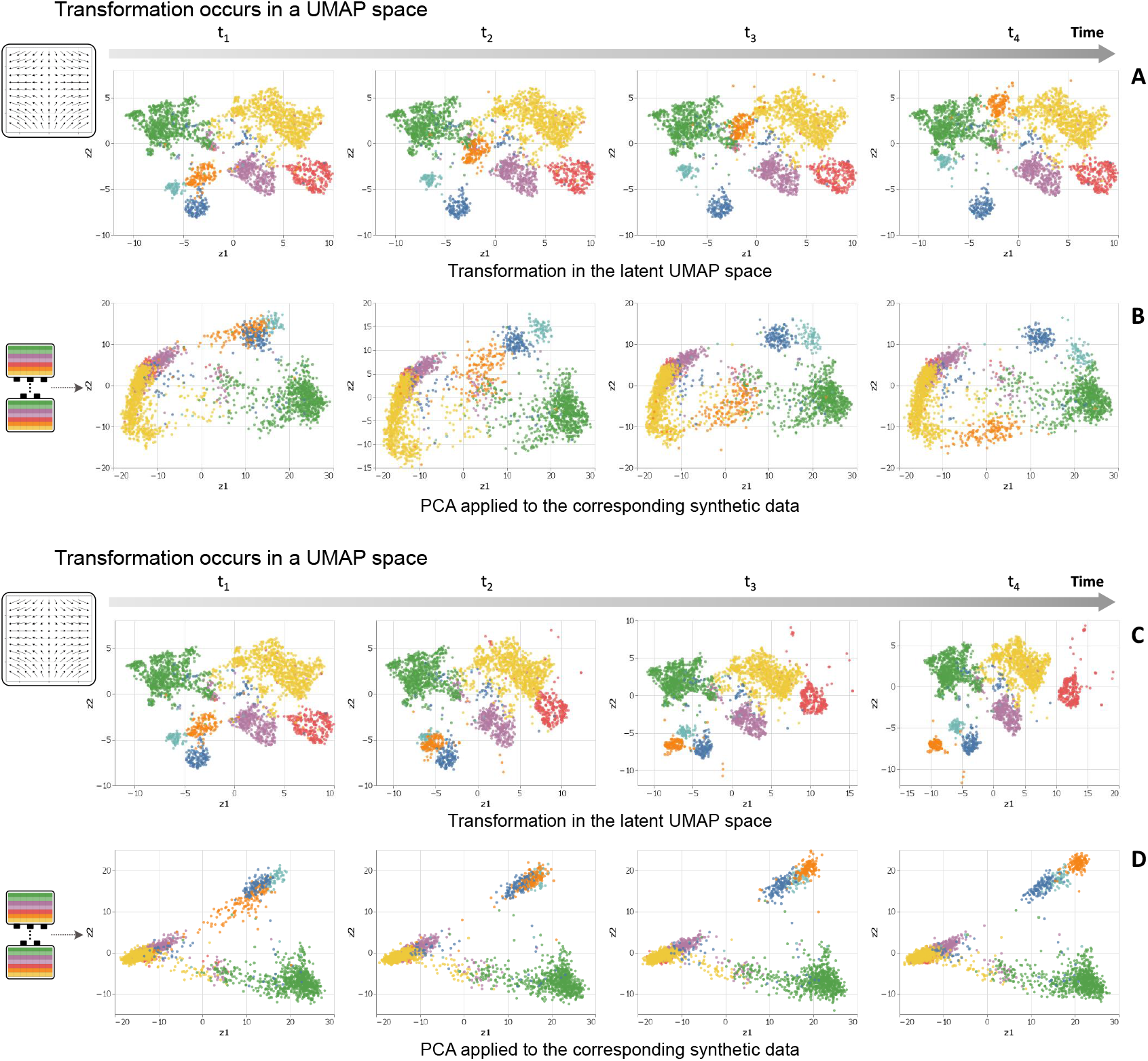
Illustrative example of a comparative analysis of how dimensionality reduction methods capture different artificially induced temporal structures within the same manifold in scRNA-seq data. **(A**,**C)**: Latent representations from the supervised VAE, trained to match the t-SNE (A) and UMAP (C) embeddings of the PBMC dataset (leftmost panel), used as initial time point (t1), and transformed to induce a visual temporal pattern corresponding to the differentiation of the blue cell type cluster (panels t2-t4). **(B**,**D)**: PCA applied to the high-dimensional datasets generated by decoding the differently transformed latent representations in the UMAP manifold using the supervised VAE. This comparison highlights how the performance of the same method (PCA) can vary depending on the transformation applied, even if this transformation is done in the same manifold.

In summary, the empirical evaluation shows that the different dimensionality reduction techniques have often different perspectives on the same underlying temporal structure and thus cannot be assumed to accurately reflect an underlying developmental process. These findings do not allow us to recommend one method as superior in general, but rather emphasize the need for careful method selection based on the characteristics of both the temporal patterns and the manifolds they exist in. In this context, our framework enables to more comprehensively characterize their performance, and can aid targeted selection of an optimal dimensionality reduction approach if some knowledge about anticipated pattern is available.

## 4 Discussion

The increasing availability of molecular datasets with complex structure and multiple measurements, such as from several time points, promises novel insights into biological mechanisms, such as understanding developmental trajectories at single-cell resolution.

For time-series scRNA-seq data, like the motivational example in Figure 1, the true underlying dynamics are typically not fully known. While there may be some knowledge about the assumed underlying process (e.g., which cell types evolve during differentiation and which markers are up- and downregulated), this is usually incomplete, and dimensionality reduction and subsequent modeling is applied precisely to infer these temporal processes. In particular, even with some knowledge of the expected driving process, it is unclear how to identify the “true” low-dimensional manifold where this process can ideally be observed. Often, researchers apply unsupervised dimensionality reduction approaches for visual inspection of temporal patterns, which cannot be guaranteed to identify this manifold, as common approaches optimize purely data-intrinsic criteria., as illustrated in Section 2.2.

To investigate to what extent different approaches can nonetheless reveal complex underlying structure such as temporal development patterns, we developed a VAE approach for generating synthetic data with temporal structure based on a chosen manifold (Figure 2). Specifically, the approach allows for defining a hypothetical “ground-truth” manifold, artificially introducing temporal structure in this space, and generating corresponding synthetic high-dimensional gene expression data.

While it was to be expected that the choice of the dimensionality reduction technique affects which patterns can be seen, comparing performance just based on visual representations is challenging for with time-series single-cell datasets, especially since there are no established benchmark datasets specifically for this purpose or ground-truth information. Our synthetic data approach provides a solution by creating such datasets based on real data that approximate an underlying dynamic system, using vector fields as an intuitive geometric framework.

We used some of these datasets to compare four popular techniques, namely PCA, t-SNE, UMAP and scVI. Although there is no quantitative measure for comparing performance – as this would, e.g., require a consistent metric for measuring distance across manifolds from different techniques – we observed in the examples that no method is consistently superior. We conclude that relying on a single techniques cannot be assumed sufficient to reliably represent dynamic patterns and thus consider it highly beneficial to look at different representations to get a more comprehensive picture. Alternatively, dimensionality reduction techniques could be extended to specifically target manifolds with the most variability over time, to flexibly pick up the manifold where the temporal dynamics actually unfold.

The main advantages of the proposed synthetic data approach are its versatility for introducing dynamic patterns via an intuitive geometric vector field approach and its flexibility with respect to different manifolds where dynamics can occur. This is enabled by a supervised component that enables the model to mimic the two-dimensional representations from different techniques. This flexibility allows for inducing different temporal patterns across various learned manifolds and posing questions such as, “We have a four-time-point dataset where one cell cluster divides over time in the manifold learned by t-SNE. Can UMAP, when trained on this generated data, also reveal this temporal pattern? Does it do so in the same way, or are there differences? Is this generalizable to any cluster division in a t-SNE space, or does it depend on the division’s speed? How significantly can parameter tuning affect the visualization?”.

When researchers have prior knowledge or hypotheses regarding the dynamics they expect, experimenting with our approach can help to identify the most suitable underlying low-dimensional manifold where the expected processes are best observed.

A limitation of the current approach is that it assumes an equal number of cells at each time point, which in reality might differ considerably. This could by addressed by applying random dropout to cells at later time points. The synthetic data approach could be further enhanced by incorporating more complex patterns. Future work could also focus on developing quantitative measures to directly compare the performance of different techniques, offering a standardized way to evaluate manifold learning in time-series single-cell data.

In summary, our approach addresses a critical gap in analyzing time-series scRNA-seq data, offering a way to reason about discrepancies between dimensionality reduction techniques. By leveraging synthetic datasets with controlled temporal dynamics, we provide a tool for exploring how different methods capture key developmental patterns, enabling researchers to assess multiple perspectives for adequately representing underlying biological processes.

## 5 Code and data availability

The exemplary time-series dataset is the *Embryoid body data*. For generating synthetic data we use two publicly available snapshot scRNA-seq datasets, the *PBMC8k data*, a dataset of peripheral blood mononuclear cell (PBMC) from [64] and the *Zeisel data*, a heterogeneous dataset of mouse brain cells [65]. Details of data acquisition and pre-processing can be found in the Supplementary Material.

We use the Julia programming language [66] of version 1.6.7 for all our analysis and models. For training scVI models in Julia and their supervised encoder adaptation for targeted generation of synthetic data, we have written a Julia version of the original scVI model from [12] based on the Python scvi-tools ecosystem [15], including pre-processing functionality based on the anndata [67] and scanpy [68] packages. As this has not been comprehensively developed in Julia before, we have created a corresponding Julia package available at https://github.com/maren-ha/scVI.jl. The complete code to reproduce our analysis and all results in this manuscript can be found at https://github.com/laia-cg/scManifoldDynamics, including tutorial Jupyter notebooks for a user-friendly introduction. Details about the hyperparameters of t-SNE and UMAP, the VAE architecture and training procedure can be found in the Supplementary Material.

## Key points

- Different dimensionality reduction techniques, such as PCA, t-SNE, UMAP, and scVI, can produce inconsistent representations of temporal patterns in time-series scRNA-seq data.
- We propose a synthetic data approach using variational autoencoders (VAEs) to introduce biologically plausible temporal patterns, for reasoning about the discrepancies of the obtained representations when subsequently applying different dimensionality reduction techniques.
- The approach enables researchers to assess the ability of various methods to capture dynamic processes, in particular highlighting the limitations of relying on a single technique.
- We provide an implementation and tutorial notebooks to guide researchers in evaluating representations and interpreting single-cell dynamics effectively.

## 7 Author contributions statement

M.H. and L.C.G. developed and performed the experiments and wrote the manuscript. R.B contributed to the writing of the manuscript. H.B. proposed the idea and supervised the project. All authors reviewed the manuscript and approved the final version.

## 8 Acknowledgments

The authors thank Martin Treppner and Moritz Hess for help with data pre-processing. This work is funded by the Deutsche Forschungsgemeinschaft (DFG, German Research Foundation) – Project-ID 322977937 – GRK 2344 (MH) and Project-ID 499552394 – SFB 1597 (HB, RB, MH).

