## Supplementary Material for "Evaluating discrepancies in dimensionality reduction for time-series single-cell RNA-sequencing data"

#### 1 Figure 3 Extended

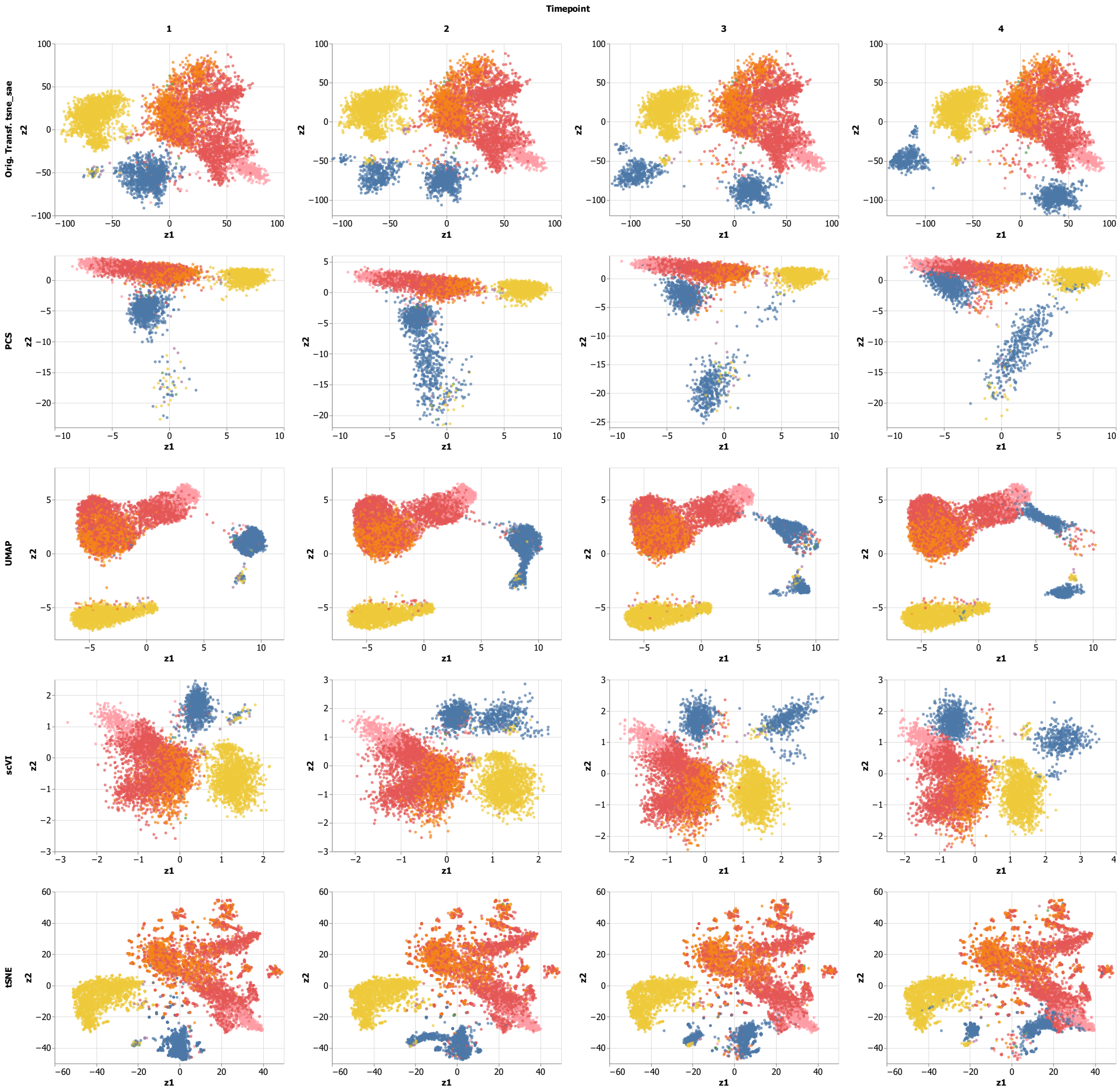


**Figure 1: Comparison of different dimension reduction approaches on scRNA-seq data with artificially introduced time structure in a t-SNE space**. **A**: Representation obtained by t-SNE on the PBMC dataset (leftmost panel), used as initial time point t1, and transformed to create synthetic time-series data (panels t2-t4), corresponding to spreading and rotation of the cell type clusters. Colors correspond to manually annotated cell types. **B-E**: Representations of the synthetic time-series dataset created from A after applying PCA (B), UMAP (C), scVI (D) and t-SNE (E).

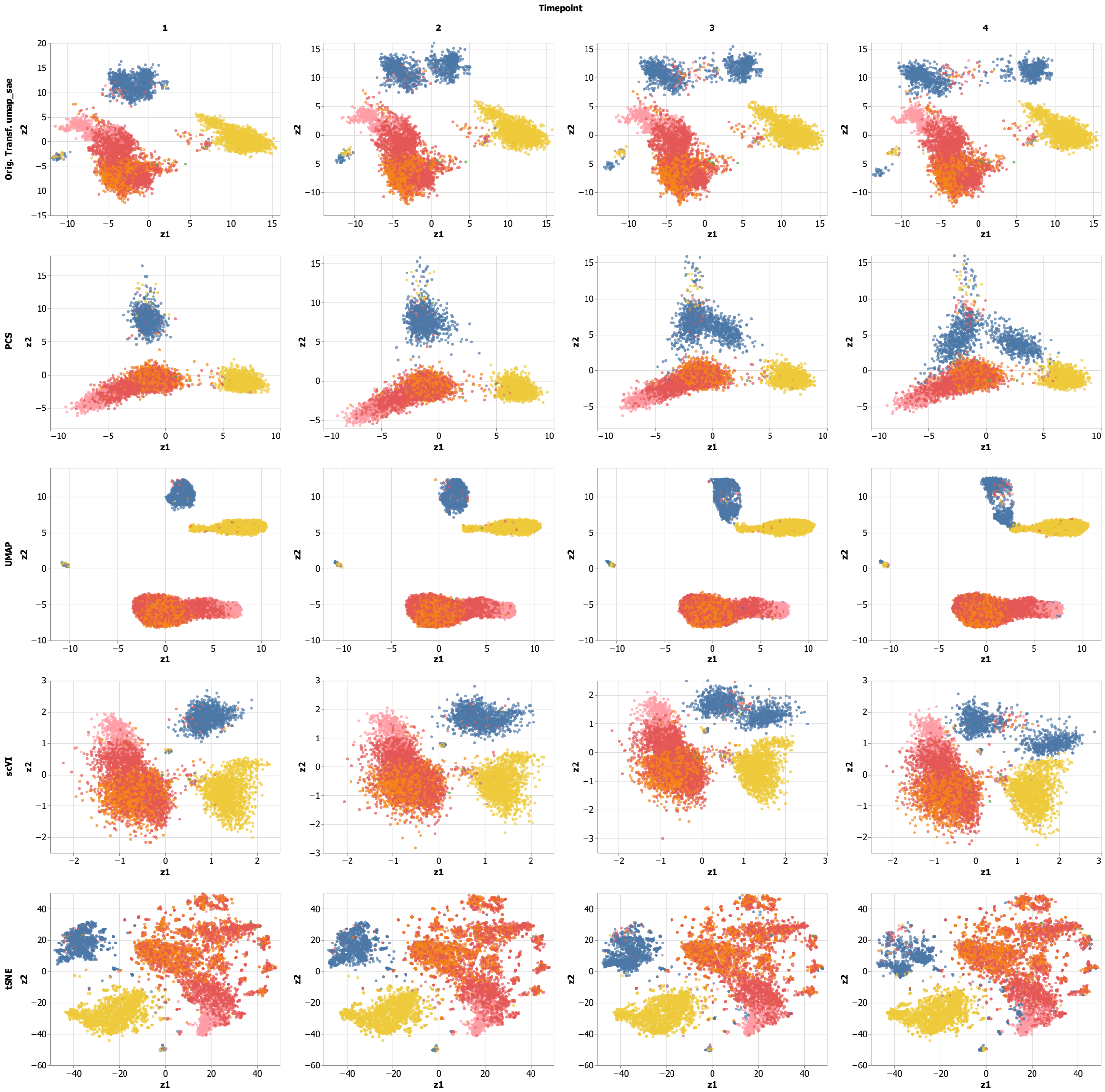


**Figure 2: Comparison of different dimension reduction approaches on scRNA-seq data with artificially introduced time structure in a UMAP space**. **A**: Representation obtained by t-SNE on the PBMC dataset (leftmost panel), used as initial time point t1, and transformed to create synthetic time-series data (panels t2-t4), corresponding to spreading and rotation of the cell type clusters. Colors correspond to manually annotated cell types. **B-E**: Representations of the synthetic time-series dataset created from A after applying PCA (B), UMAP (C), scVI (D) and t-SNE (E).

#### 2 Figure 4 Extended

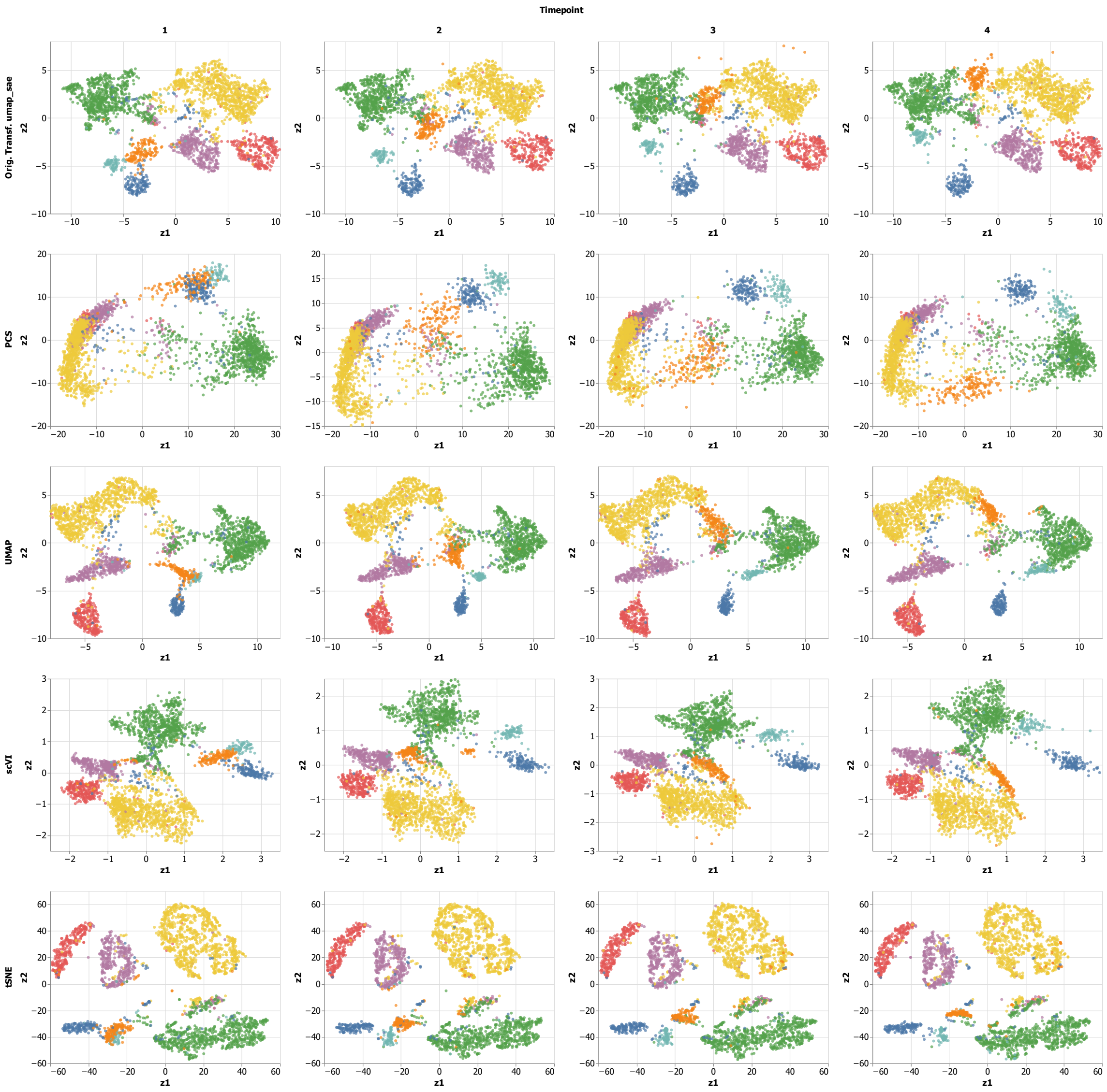


**Figure 3: Comparison of different dimension reduction approaches on scRNA-seq data with artificially introduced time structure in a UMAP space**. **A**: Representation obtained by t-SNE on the PBMC dataset (leftmost panel), used as initial time point t1, and transformed to create synthetic time-series data (panels t2-t4), corresponding to spreading and rotation of the cell type clusters. Colors correspond to manually annotated cell types. **B-E**: Representations of the synthetic time-series dataset created from A after applying PCA (B), UMAP (C), scVI (D) and t-SNE (E).

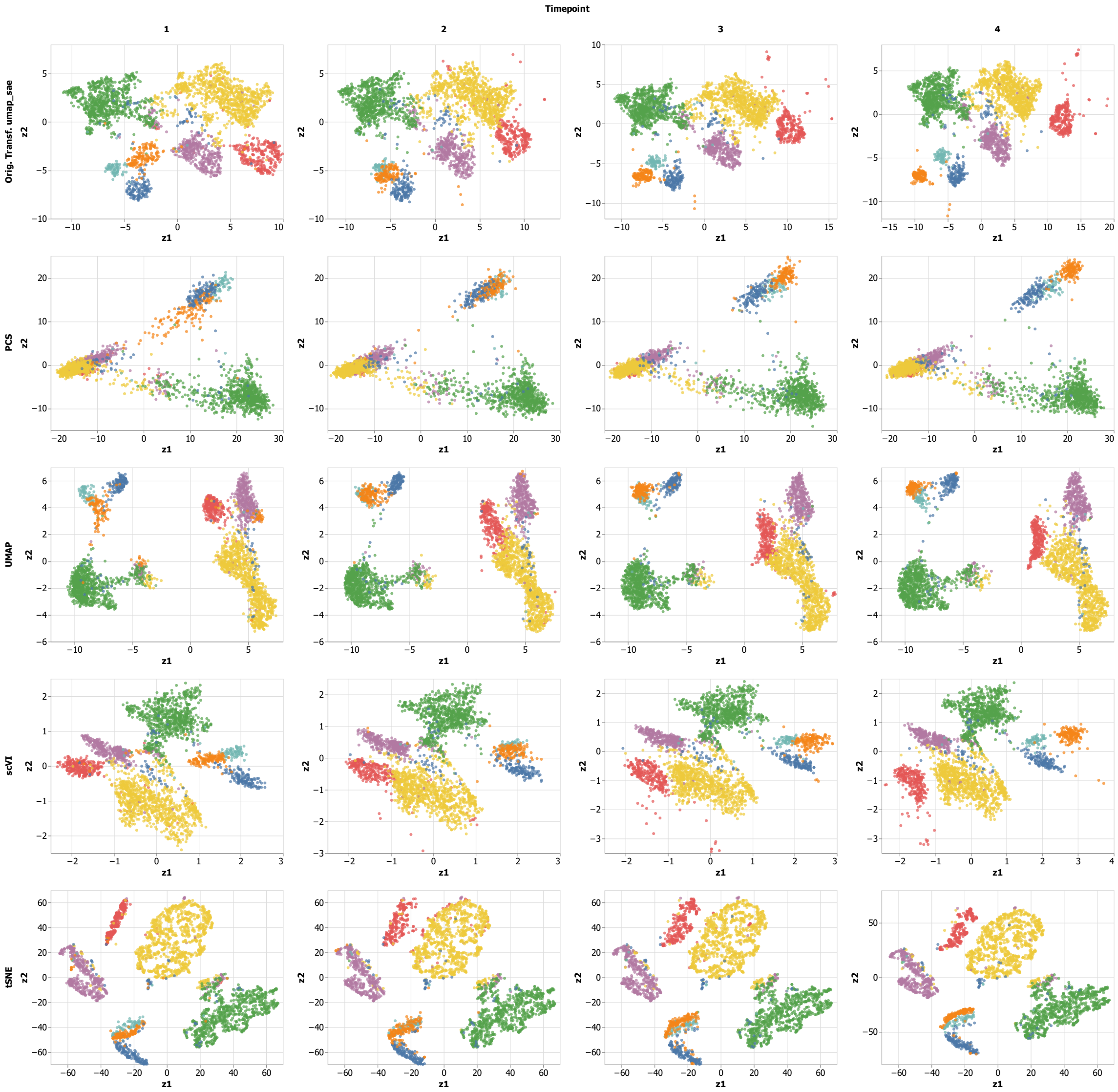


**Figure 4: Comparison of different dimension reduction approaches on scRNA-seq data with artificially introduced time structure in a UMAP space**. **A**: Representation obtained by t-SNE on the PBMC dataset (leftmost panel), used as initial time point t1, and transformed to create synthetic time-series data (panels t2-t4), corresponding to spreading and rotation of the cell type clusters. Colors correspond to manually annotated cell types. **B-E**: Representations of the synthetic time-series dataset created from A after applying PCA (B), UMAP (C), scVI (D) and t-SNE (E).

### 3 Details on pre-processing of the data

#### 3.1 Embryoid body data

The Embryoid Body dataset was obtained from Mendeley Data at https://data.mendeley.com/datasets/v6n743h5ng/1 and preprocessed according to the authors' indications described in their tutorial notebook (https://colab.research.google.com/github/KrishnaswamyLab/PHATE/blob/master/Python/tutorial/EmbryoidBody.ipynb#scrollTo=ZAdnUiMDFdEf). After pre-processing, 16821 cells remained in the dataset. For this study, 1000 highly variable genes were selected using the appropriate function available in the scVI Julia package.

#### 3.2 PBMC data

The PBMC dataset was obtained from the 10x Genomics platform (8k PBMCs from a Healthy Donor) and processed according to Bioconductor workflows [1]. After pre-processing, 7820 cells remained in the dataset. The specific R script used for pre-processing is available in the provided repository. For this study, 200 highly variable genes were selected using the same methodology used in the referenced workflow.

#### 3.3 Zeisel data

The Zeisel dataset was obtained from the R package `scRNAseq` under the function `ZeiselBrainData()`. It is also available under the GEO accession code GSE60361 and was preprocessed following the workflow described in the Bioconductor online book (http://bioconductor.org/books/3.14/OSCA.workflows/zeisel-mouse-brain-strt-seq.html). This workflow is part of the larger study *Orchestrating single-cell analysis with Bioconductor* [1]. After pre-processing, 2816 cells remained in the dataset. For this study, 1000 highly variable genes were selected using the same methodology used in the referenced workflow.

### 4 Details on the hyperparameter selection for the dimensionality reduction approaches

#### 4.1 Embryoid Body data

- Snapshot dataset
  - PCA was performed on log-normalized counts.
  - t-SNE was performed on the first 50 principal components, with a perplexity value of 20. All other hyperparameters were set to their default values as specified in the t-SNE Julia package (https://github.com/lejon/t-SNE.jl).
  - UMAP was performed on log-normalized counts.

#### 4.2 PBMC data

- Snapshot dataset
  - PCA was performed on log-normalized counts.
  - t-SNE was performed on the first 35 principal components, with a perplexity value of 50. All other hyperparameters were set to their default values as specified in the t-SNE Julia package.

- UMAP was performed on the first 35 principal components, with a `min_dist` value of 0.5. All other hyperparameters were set to their default values as specified in the UMAP Julia package.

- Synthetic time-series dataset
  - PCA was performed on log-normalized counts.
  - t-SNE was performed on the first 20 principal components, with a perplexity value of 200. All other hyperparameters were set to their default values as specified in the t-SNE Julia package.
  - UMAP was performed on the first 20 principal components. All other hyperparameters were set to their default values as specified in the UMAP Julia package.

##### 4.3 Zeisel data

- Snapshot dataset
  - PCA was performed on log-normalized counts.
  - t-SNE was performed on the first 100 principal components, with a perplexity value of 20. All other hyperparameters were set to their default values as specified in the t-SNE Julia package.
  - UMAP was performed on the first 100 principal components, with a `min_dist` value of 0.5. All other hyperparameters were set to their default values as specified in the UMAP Julia package.

- Synthetic time-series dataset
  - PCA was performed on log-normalized counts.
  - t-SNE was performed on the first 20 principal components, with a perplexity value of 200. All other hyperparameters were set to their default values as specified in the t-SNE Julia package.
  - UMAP was performed on the first 20 principal components. All other hyperparameters were set to their default values as specified in the UMAP Julia package.

#### 5 Details on the training parameters of scVI

The version of scVI used in this project is the Julia version of scVI, available at https://github.com/maren-ha/scVI.jl. The model was always trained on raw counts with a batch size of 128 and for a maximum of 200 epochs. The optimizer used was ADAM with decay regularization, specifically, with a learning rate of $1e-3$ and a weight decay parameter of $1e-6$. All other training parameters were set to their default values as specified in the scVI Julia package.
